# Anorectal care products cause variable inflammation and adverse barrier effects in a mucosal epithelial model

**DOI:** 10.64898/2026.10.30.763242

**Authors:** Erik C Swanson, Christopher M Basting, Ty Schroeder, Greg Lam, Evan Goldstein, Nichole R Klatt

**Author notes:** **Corresponding Author** Nichole R. Klatt –.

## Abstract

**Introduction:** Sexually transmitted infections (STIs) and colorectal cancer (CRC) are rising in the United States, yet their shared mucosal environment remains underexplored. Rectal douching, a common practice among individuals engaging in receptive anal intercourse, has been epidemiologically linked to increased STI and HPV risk, but biological mechanisms are unclear.

**Methods:** Using a human colorectal epithelial co-culture model (Caco-2/HT29-MTX-E12), we compared the effects of common douching products on cell viability, barrier integrity, and cytokine response.

**Results:** Hyperosmolar or surfactant-containing solutions, including saline enemas, detergents, and soaps, caused rapid declines in metabolic activity and transepithelial electrical resistance, coupled with IL-6 and TNF-α upregulation. In contrast, isotonic formulations maintained epithelial viability and barrier function comparable to buffered controls while reducing inflammatory signaling.

**Discussion:** These findings provide mechanistic evidence that many consumer douching products compromise mucosal health and highlight the feasibility of harm-reduction designs for safer anal hygiene and sexual preparation.

## Introduction

Sexually transmitted infections (STIs) in the United States have resurged after years of progress, with recent national estimates of 67 million concurrent cases and 26 million annual incidental cases, a healthcare burden of approximately $16 billion (2018 USD) in associated medical costs^1–5^. In parallel, colorectal cancer (CRC) rates, particularly early-onset CRC, have increased over the past decade, a trend documented in population-based cancer surveillance and contemporary cohort analyses^6–8^. These two trajectories are usually considered separately: STIs as transmissible mucosal infections concentrated in sexual networks, and CRC as a multifactorial malignancy influenced by age, genetics, inflammation, microbiota, and environmental exposures. Yet both disease areas intersect at the anorectal mucosa, where sexual behaviors, local products, and intimate care practices may alter epithelial integrity, susceptibility to infection, and possibly long-term carcinogenic pathways.

Receptive anal intercourse (RAI) is common across multiple populations including men who have sex with men (MSM) but also among women and heterosexual men. National survey studies have reported substantial lifetime experience of anal sex among U.S. adults, including women and younger adults^9,10^. More focused studies in Brazil, France, Europe, and the United States consistently show high RAI prevalence among MSM and other individuals at high risk of HIV^9,11–16^. Within these groups, rectal douching and enema use before RAI is widely practiced. Data suggest that a large majority of people who engage in RAI douche at least occasionally^11–14^. This makes douching one of the most prevalent, modifiable anorectal exposures in sexually active populations.

A growing body of epidemiologic work links rectal douching to higher odds of rectal STIs and other anorectal morbidity. Among MSM using HIV pre-exposure prophylaxis, douching has been associated with rectal gonorrhea and chlamydia, even after accounting for sexual risk behaviors^13^. Similar associations have been described for lymphogranuloma venereum (LGV) proctitis in MSM, where anal enema use clustered with other high-risk practices^17^. A systematic review and meta-analysis reported that douching was associated with 2.5–3.25-fold higher odds of HIV, HPV, and other STIs, suggesting that the practice may amplify acquisition risk beyond behavioral confounding^18,19^. Mechanistically, this is plausible: hyperosmolar or otherwise irritating products can damage colorectal epithelium, increase permeability, and induce local inflammation, which in turn could enhance susceptibility to viral or bacterial pathogens^20–22^.This pattern mirrors the more mature literature on vaginal douching, which has been tied to adverse reproductive outcomes, increased HPV infection, and cervical neoplasia, leading to longstanding recommendations against routine douching^1,23–25^.

Despite this, rectal douching products and routines have not been systematically evaluated for their effects on the human anorectal barrier. Most available data comes from microbicide-enema development studies showing that iso-osmolar formulations are safer and better distributed than hyper- or hypo-osmolar preparations, implying that current consumer products may span a wide and largely unregulated spectrum of epithelial toxicity^21,22^. Importantly, existing STI and CRC surveillance reports do not incorporate douching as an exposure, so no comprehensive estimate of douching-attributable health burden exists^1–3,6,8^ Yet if douching even modestly increases HPV, HIV, and bacterial STI incidence in high-prevalence networks, the downstream costs could be substantial, given that STIs and HPV-related disease together generate billions of dollars in direct medical expenditures annually^2,4^. This makes douching a strategically attractive, but currently overlooked, point of intervention that sits at the junction of infectious-disease prevention and cancer prevention.

Our study was designed to begin filling this gap. We developed a simple epithelial model that recapitulates key features of the human anorectal surface and used it to compare commonly used douching solutions. We quantified effects on epithelial integrity, cell viability, and inflammatory response, thereby providing a biologically coherent link between a prevalent behavior and previously observed increases in anorectal STI/HPV risk. In doing so, we identify specific formulations with markedly lower mucosal impact, supporting the feasibility of a harm-reduction approach to intimate hygiene and anal-sex preparation. Collectively, these data should prompt more explicit consideration of douching in STI and CRC prevention frameworks and justify more nuanced in vivo and clinical studies to refine safe-use recommendations

## Results

### Metabolic Activity and Cellular Viability

Resazurin assays revealed striking differences in epithelial metabolic activity among treatment conditions (Figure 1). Exposure to common douching solutions or additives, including Fleet, detergent (SDS), hand soap, caused sharp and sustained reductions in metabolic activity relative to DMEM control (adjusted *p* < 0.001 for all contrasts at 72 h). These decreases were evident within the first hour post-exposure and expanded over 72-hour observation window, indicating cytotoxicity rather than transient metabolic suppression. In contrast, Future Method Powder (FMP) and phosphate-buffered saline (PBS) maintained metabolic activity levels statistically indistinguishable from DMEM across all time points (adjusted *p* > 0.05). Tap water, deionized water, and ethanol produced mild to moderate metabolic impairment, consistent with osmotic and solvent stress effects. Numeric results are summarized in Table 1. Together, these data suggest that several widely used consumer formulations or product additives rapidly compromise epithelial cell viability, whereas the powder formulation of Future Method preserves metabolic integrity comparable to isotonic buffered controls.

**Table 1.** Summary of endpoint TEER and Resazurin responses relative to DMEM control. Mean ± standard error (SE) and Benjamini–Hochberg (BH)–adjusted *p*-values at the final measurement timepoint for each condition are shown. TEER reflects epithelial barrier integrity (fold-change relative to baseline), and Resazurin (inverse Ab_605_) reflects metabolic activity/viability. Conditions are ordered approximately by similarity to DMEM (Dulbecco’s Modified Eagle Medium, control). Missing values indicate conditions or endpoints not measured at that timepoint.

|  | TEER (mean ± SE) | Resazurin (mean ± SE) | TEER p (BH) | Resazurin p (BH) | TEER final t | Rez final t |
|---|---|---|---|---|---|---|
| **DMEM** | 0.92 ± 0.07 | 2.57 ± 0.29 | NA | NA | 24.0 | 72.0 |
| FMP | 0.99 ± 0.02 | 2.82 ± 0.16 | 0.956 | 0.825 | 24.0 | 72.0 |
| PBS | 1.02 ± 0.03 | NA | 0.956 | NA | 24.0 | NA |
| NaCl | NA | 2.35 ± 0.25 | NA | 0.825 | NA | 72.0 |
| diH2O | 0.43 ± 0.28 | 2.17 ± 0.12 | 0.005 | 0.459 | 24.0 | 72.0 |
| H2O | 0.18 ± 0.00 | 2.16 ± 0.07 | <0.001 | 0.459 | 24.0 | 72.0 |
| SEVE | 0.15 ± 0.03 | NA | <0.001 | NA | 24.0 | NA |
| FLEET | 0.14 ± 0.02 | 1.05 ± 0.04 | <0.001 | <0.001 | 24.0 | 72.0 |
| SDS | 0.12 ± 0.02 | 1.20 ± 0.03 | <0.001 | <0.001 | 24.0 | 72.0 |
| NA | NA | 2.04 ± 0.15 | NA | 0.231 | NA | 72.0 |
| NA | NA | 1.14 ± 0.03 | NA | <0.001 | NA | 72.0 |

**Figure 1.**
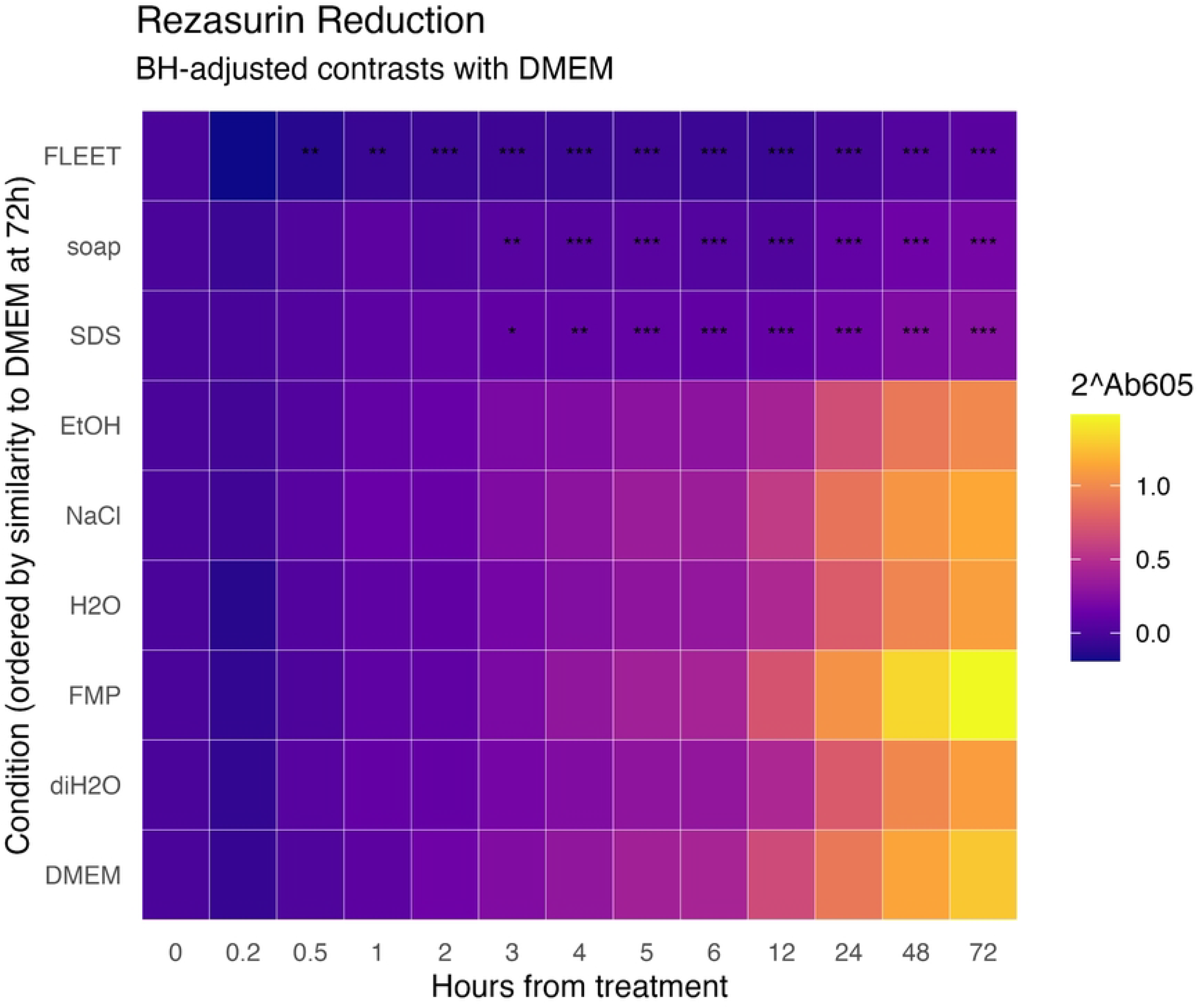
Heatmap of metabolic activity (inverse Ab605) over time relative to DMEM control. Metabolic activity of Caco-2/HT29-MTX-E12 (10:1) co-cultures was quantified by resazurin reduction (Absorbance_605_ nm) over 72 hours following exposure to each condition. Values were converted to 2^Ab605, proportional to relative metabolic activity, and normalized within replicates. Colors represent the mean inverse absorbance across replicates (yellow = higher, blue = lower). Missing values were imputed by linear interpolation, and rows are ordered by final similarity to DMEM (Dulbecco’s Modified Eagle Medium, control). Conditions include FMP (Future Method Powder), SDS (sodium dodecyl sulfate detergent), FLEET (saline enema solution), SOAP (dilute hand soap), NaCl (saline), H_2_O (tap water), diH_2_O (deionized water), and EtOH (ethanol). Asterisks denote Benjamini–Hochberg–adjusted *p*-values for pairwise contrasts with DMEM (*p* < 0.05; ^*^*p* < 0.01; ^**^*p* < 0.001). (N=6 in each case)

### Epithelial Barrier Integrity

Measurements of transepithelial electrical resistance (TEER) paralleled the metabolic results and provided quantitative insight into barrier disruption dynamics (Figure 2). Barrier integrity declined dramatically following exposure to SDS, soap, Fleet, and Summer’s Eve, with TEER values falling below 0.2-fold baseline within 24 h (*p* < 0.001 vs DMEM). These losses persisted through the recovery phase, indicating incomplete reconstitution of tight-junction function. Conversely, FMP, PBS, and saline (NaCl) preserved TEER near baseline values (0.9–1.1-fold), suggesting negligible impact on epithelial junctional complexes. Deionized and tap water caused modest, transient decreases in TEER, likely reflecting osmotic stress that partially resolved within 48 h. The close correspondence between metabolic impairment and TEER loss across conditions highlights a common underlying mechanism of membrane or junctional injury. As in resazurin assays, FMP demonstrated the least epithelial disruption.

**Figure 2.**
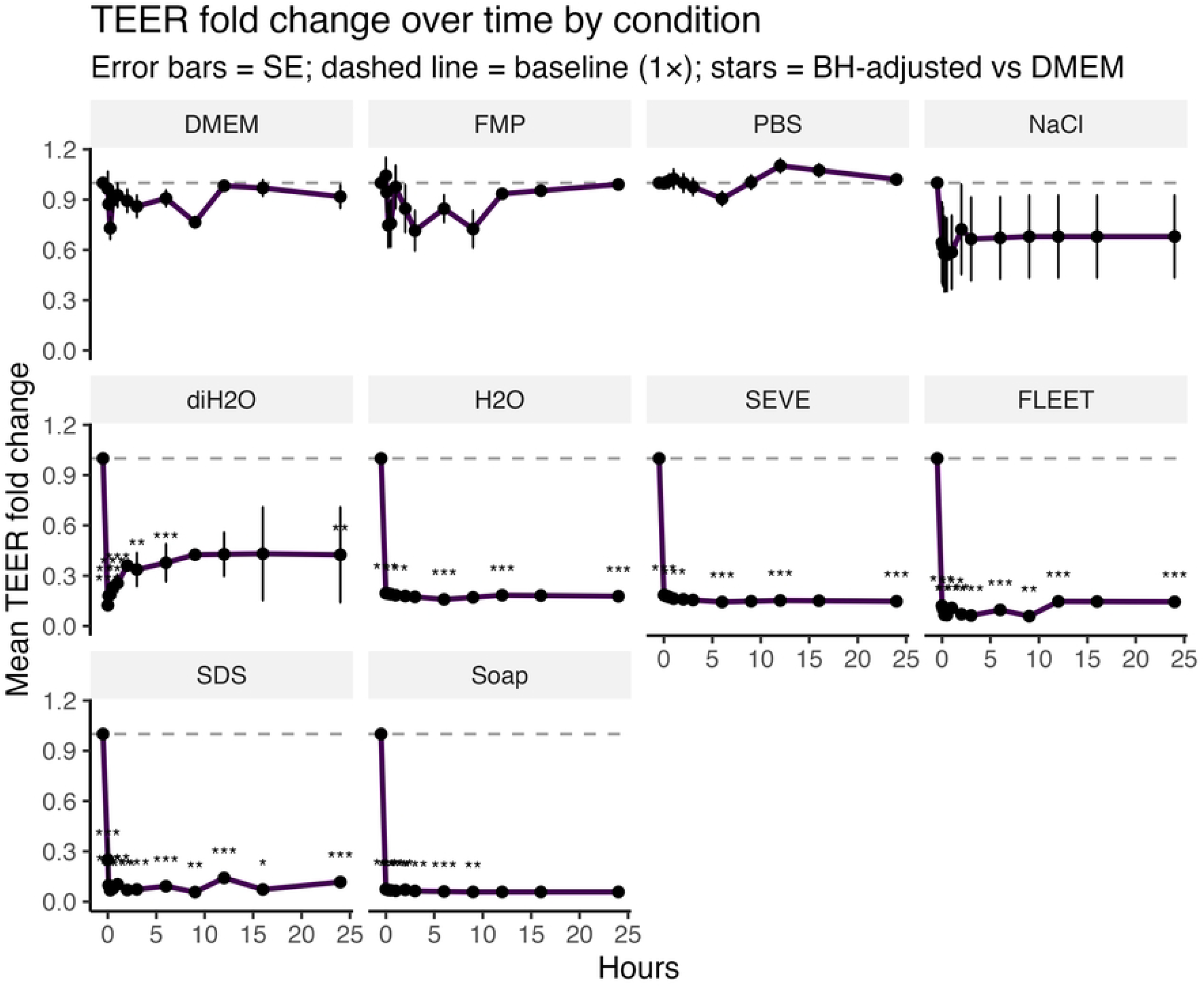
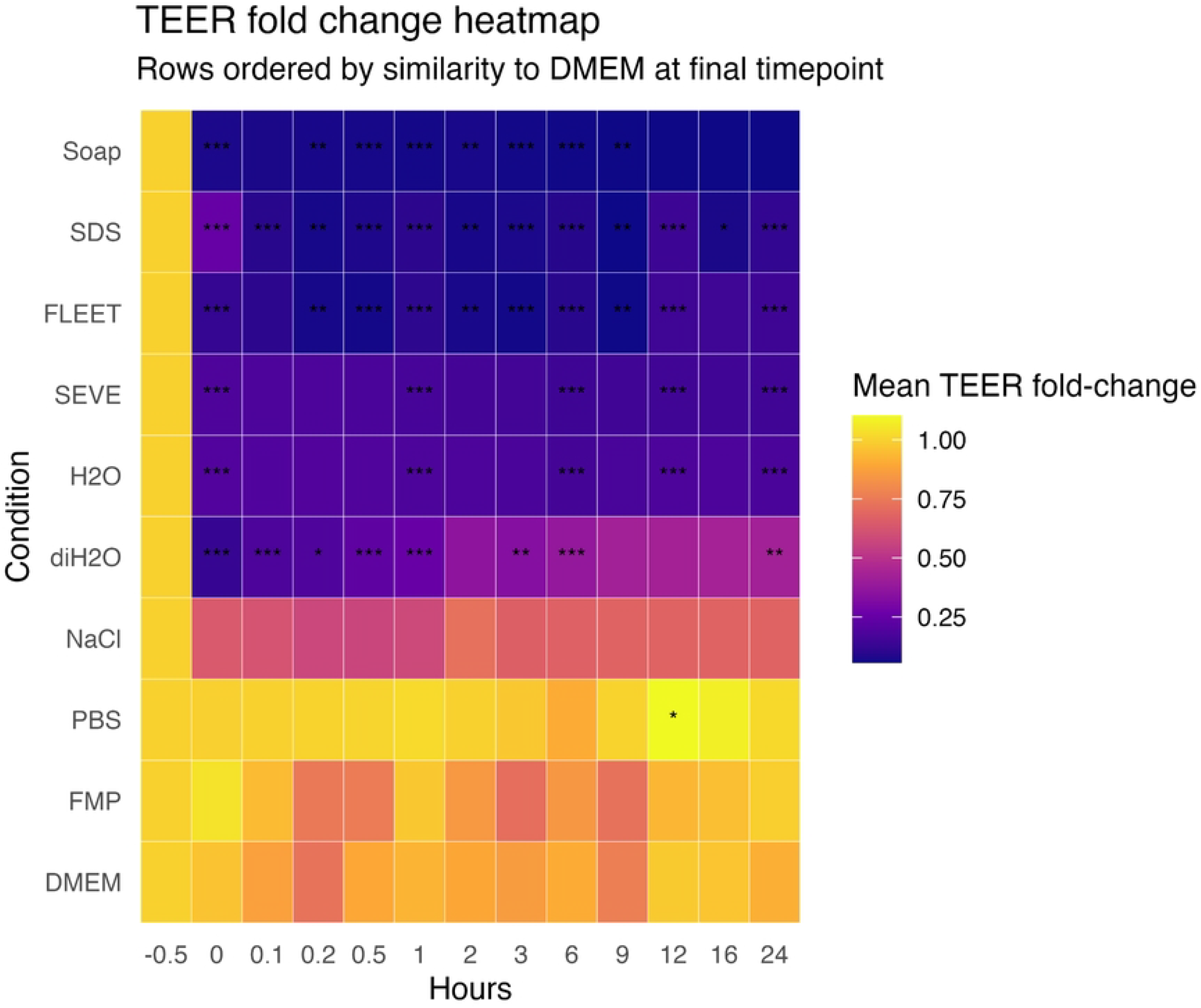
Fold change in transepithelial electrical resistance (TEER) in Caco-2/HT29-MTX-E12 10:1 co-cultures following exposure. TEER was measured over time following exposure to each condition and normalized to the pretreatment (–0.5 h) baseline for each replicate (fold change = TEER / baseline TEER). **A)**.Lines show mean ± standard error (SE) of normalized TEER across replicates; the dashed gray line marks baseline TEER (1x fold change). **B)**. Colors indicate the mean normalized TEER (yellow = higher, blue = lower) across all replicates per condition and timepoint, with missing values imputed by linear interpolation. Rows are ordered by final similarity to DMEM. Conditions: DMEM (Dulbecco’s Modified Eagle Medium, control), FMP (Future Method Powder), PBS (phosphate-buffered saline), NaCl (saline), diH_2_O (deionized water), H_2_O (tap water), SEVE (*Summer’s Eve* feminine wash), FLEET (saline enema solution), SDS (sodium dodecyl sulfate) and Soap (dilute hand soap). Asterisks denote Benjamini–Hochberg–adjusted p-values for pairwise contrasts with DMEM (*p* < 0.05; \**p* < 0.01; \*\**p* < 0.001). Each facet represents one condition’s mean TEER trajectory over time. (N=6-9)

### Inflammatory Cytokine Response

Cytokine quantification revealed a distinct inflammatory signature associated with douching agents (Figure 3). Fleet and SDS elicited marked up-regulation of IL-6 (log_2_ fold-change > 4; adjusted *p* < 0.01) and moderate induction of TNF-α (log_2_ fold-change ***, adjusted *p* < ***), consistent with epithelial stress and pro-inflammatory signaling. Even ostensibly mild perturbations such as tap water triggered significant IL-6 release, underscoring the sensitivity of the rectal mucosal model to hypotonic or non-buffered exposures. In contrast, FMP not only failed to induce cytokine expression but significantly reduced IL-6 relative to baseline (*p* < 0.05 for IL-6 down-regulation), suggesting potential anti-inflammatory or protective properties of the formulation. Soap produced an intermediate response, reflecting partial epithelial activation relative to SDS or Fleet. Collectively, these findings indicate that epithelial inflammation parallels cytotoxicity and barrier compromise, and that hypotonicity and surfactant content are key determinants of this response.

**Figure 3.**
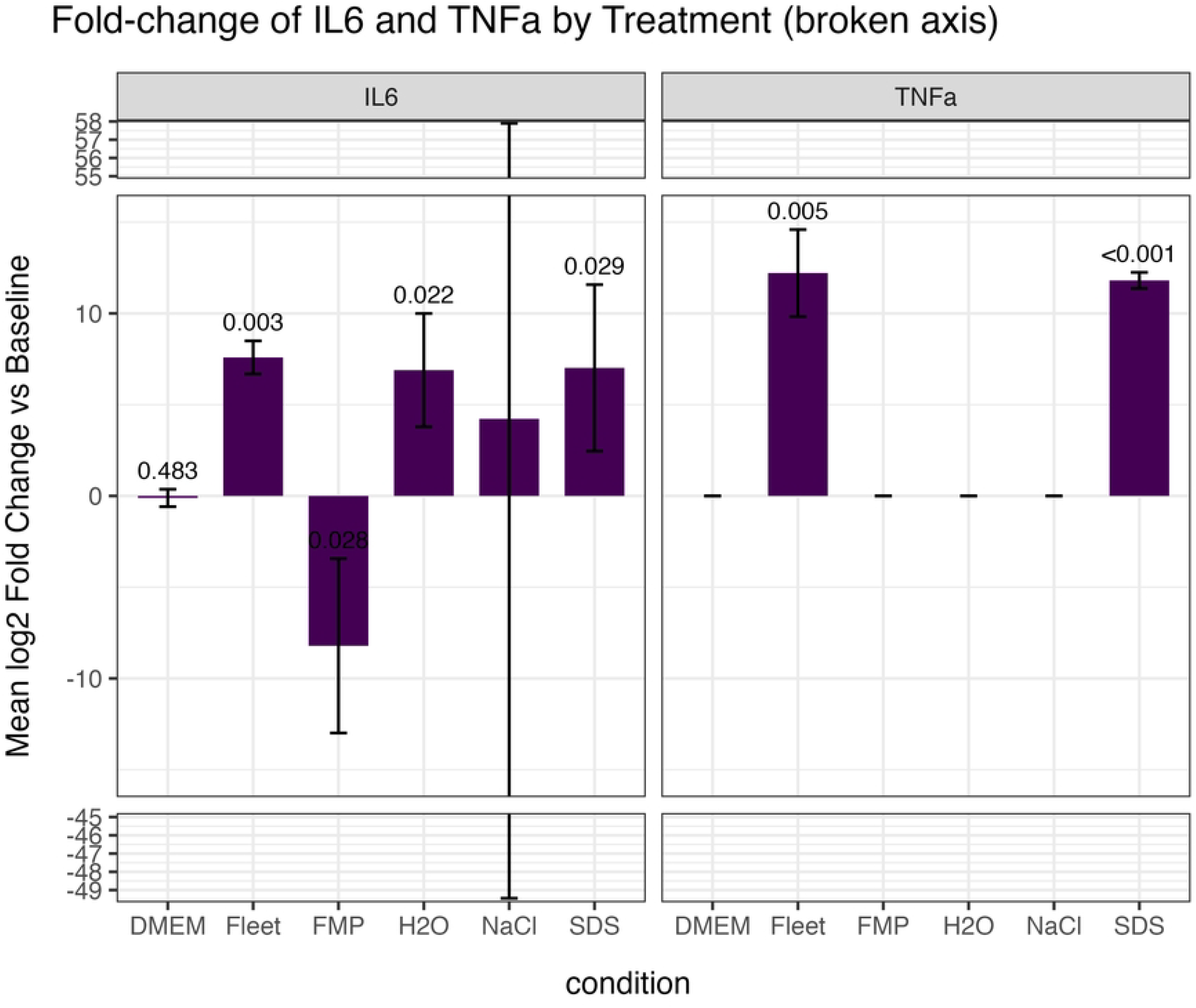
Fold-change in IL-6 and TNFα expression following treatment. Cytokine concentrations in Caco-2/HT29-MTX-E12 (10:1) co-cultures were measured 24 hours after treatment and expressed as log_2_ fold-change relative to baseline (pretreatment) levels. Bars represent mean ± standard error (SE) across biological replicates. Conditions include DMEM (Dulbecco’s Modified Eagle Medium, control), FMP (Future Method Powder), H_2_O (tap water), NaCl (saline), SDS (sodium dodecyl sulfate detergent), and FLEET (saline enema solution). *p*-values are Benjamini–Hochberg–adjusted one-sample *t*-tests comparing mean log_2_ fold-change to zero (no change). The broken y-axis emphasizes large positive and negative expression shifts while preserving relative differences among treatments. (N=3 in each case)

**Figure S1.**
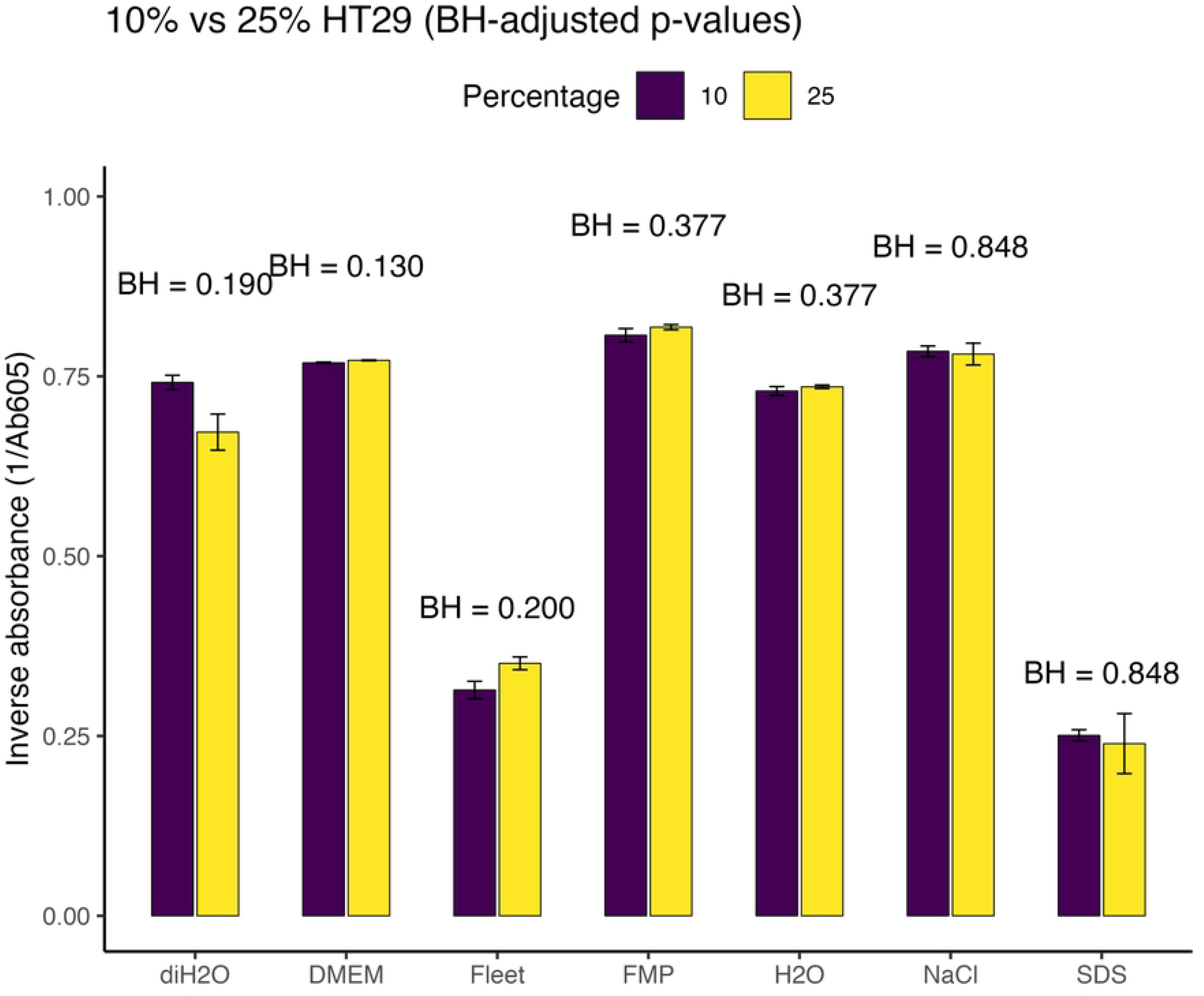

### Integrated Summary of Cellular Outcomes

Table 1 summarizes final-timepoint outcomes for metabolic activity and TEER. Across endpoints, FMP consistently clustered with DMEM and PBS controls, while SDS and Fleet ranked among the most disruptive conditions. The data collectively identify FMP as a low-toxicity, low-inflammation alternative to conventional douching solutions, whereas several widely marketed products and tap water compromise epithelial homeostasis within minutes of contact.

## Discussion

This study provides new physiological insights about consumer douching products in a controlled epithelial model, revealing significant impacts on mucosal integrity, viability, and inflammatory response. Prior epidemiologic work has associated rectal douching with elevated odds of HIV, HPV, and bacterial STIs among men who have sex with men and other populations^13,18,19^, but direct mechanistic evidence has been lacking. Our findings extend that limited data by showing that widely used douches including saline enemas, detergents, and soaps rapidly impair epithelial barrier function and induce pro-inflammatory signaling, providing a plausible biological basis for observed epidemiologic associations. In contrast, an isotonic, powder-based formulation (FMP) preserved metabolic activity, TEER, and cytokine homeostasis comparable to buffered controls, supporting the feasibility of harm-reduction–oriented product design.

These results align with limited mucosal barrier research emphasizing the importance of osmolarity and surfactant content in determining epithelial safety. Studies of rectal and vaginal microbicide vehicles have shown that hyperosmolar lubricants cause epithelial damage and increase pathogen susceptibility, while iso-osmolar formulations show decreased barrier effects^20,21^. Similarly, the vaginal douching literature has long linked mucosal disruption and altered microbiota to increased risk of HPV infection and neoplasia^23^. By demonstrating that even brief exposure to unbuffered or hyperosmolar solutions can compromise junctional integrity, our data underscore how epithelial perturbation may act as an upstream driver of infection risk and possibly inflammation-mediated oncogenic pathways.

Despite its utility, our co-culture model remains a simplified in-vitro system lacking complex tissue structure, immune cells, and microbiota that shape mucosal resilience in vivo. Such models cannot capture restitution kinetics, microbiota-derived protection, or the influence of repeated exposures. Future studies should extend these findings using organoid, ex-vivo, or clinical approaches to validate epithelial thresholds and long-term safety.

Limitations notwithstanding, the dramatic difference between treatments indicates strong effect. Until rigorous safety data become available, our results argue for greater caution in the use of unregulated and untested douching products and for clearer safety standards based on osmolarity, pH, and surfactant content. Incorporating these considerations into STI and colorectal-cancer prevention frameworks may represent a pragmatic, low-cost opportunity to mitigate mucosal injury and its downstream infectious and inflammatory sequelae.

## Methods

### Cell lines and maintenance

Two human colorectal epithelial cell line were used in this study. Caco-2 and HT29-MTX-E12 cells were maintained under sterile conditions in DMEM supplemented with 10% fetal bovine serum (FBS), 1× penicillin–streptomycin (anti-anti), and 1× non-essential amino acids (NEAA). Media were prepared fresh as needed to reduce contamination risk. Individual cultures were initiated from low passage cryopreserved stocks stored in liquid nitrogen. Cells were thawed rapidly in a 37 °C water and diluted in warm medium, pelleted (≈700–1,300 × g, 7–10 min), resuspended, and seeded to T-25 flasks. Passaging was performed at ∼80% confluence using 0.25% trypsin-EDTA (3–5 min at 37 °C), followed by neutralization in complete DMEM, gentle trituration to minimize clumping, centrifugation, and reseeding. For cryopreservation, cell suspensions with ≥90% viability were frozen at 2–2.5 × 10^6 cells/mL in 90% FBS/10% DMSO, held at −80 °C overnight, and transferred to liquid nitrogen for long-term storage.

### Co-culture model and seeding

Caco-2:HT29-MTX-E12 co-cultures were established at a 10:1 ratio based on prior literature supporting their use as an intestinal mucosal model. For plate-based viability assays, cells were seeded at 2 × 10^5^ cells/cm^2^ and maintained for ∼14 days with medium changes every 2–3 days. For barrier assays, polyester transwell inserts (growth area 1.12 cm^2^, pore size 0.4μm) were seeded at a total of 2.0 × 10^5^ cells per insert (10:1 Caco-2:HT29-MTX-E12) and cultured with 1.5 mL DMEM basolaterally and 0.5 mL apically. Cultures were maintained for ∼14 days prior to experimental exposures, based on our pilot measurements indicating that TEER exceeded 200 Ω within a few days and reached ∼1,500 Ω within 2 weeks.

### Test solutions and preparation

Test conditions comprised: DMEM (control), tap water (H_2_O; 0.22 µm filtered), deionized water (diH_2_O; 0.22 µm filtered), saline made in deionized water (NaCl; 10 g/L; 0.22 µm filtered), PBS (pH 7.4), FLEET® saline enema (with mono and dibasic sodium phosphate, 7/19 grams per 250mL), sodium dodecyl sulfate (SDS; 1% w/w in deionized water; 0.22 µm filtered), liquid hand soap (“SoftSoap”; 1% w/w in tap water; 0.22 µm filtered), ethanol (EtOH; 1% v/v in tap water; 0.22 µm filtered), and Future Method formulation (FMP, powdered product diluted in tap water; 0.22 µm filtered).

### Exposure regimen

Immediately prior to exposure, baseline measurements (TEER for transwells; absorbance for viability plates) were recorded. Apical medium was aspirated and replaced with 500 µL of the assigned treatment for 5 min. Following exposure, apical chambers were rinsed and refilled with 500 µL DMEM and the basolateral chamber with 1.5 mL DMEM. Sampling and measurements were taken for 9-72 hours based on preliminary data.

### Transepithelial electrical resistance (TEER)

TEER was measured on transwells beginning at culture day ∼5 and at all exposure time points using standard chopstick electrodes. For analysis, replicate-level TEER values were normalized to each replicate’s pre-treatment baseline to yield fold change. Condition summaries (mean ± SE) were generated at each time point. Per-timepoint differences from DMEM were estimated by fitting, at each time, a linear model (TEER ∼ condition) and computing estimated marginal means (EMMs) with contrasts versus DMEM; p-values were adjusted across times/conditions using the Benjamini–Hochberg (BH) method. For visualization, missing condition means at intermediate times were linearly interpolated within condition (zoo::na.approx with rule = 2) to facilitate continuous heatmaps/trajectories; all inference used observed (non-imputed) data.

### Metabolic activity/viability (resazurin reduction)

Co-cultures in tissue-treated 48-well plates were exposed as above. Resazurin working solution was added immediately post-exposure, and absorbance at 605 nm (Ab605) was recorded at baseline, 15, 30, and 60 min, hourly to 8 h, and then periodically to 72 h. To obtain a metric proportional to inverse absorbance (higher values = higher metabolic activity in our plots), Ab605 was transformed as 2^Ab605 (“inverse Ab605”) and summarized over time (mean ± SE). Condition effects over time were modeled with a linear mixed model of the transformed outcome: *invAb ∼ condition × ns(time, df = 4) + (1* | *replicate)*, followed by marginal means (EMMeans) averaged over the observed time distribution (weights = “proportional”) and pairwise contrasts among conditions with BH correction. For time-resolved effect displays relative to DMEM, we also fit per-time linear models (Ab605 ∼ condition) and reported BH-adjusted contrasts.

### Cytokine measurements (IL-6 and TNF-α)

Apical media were collected from selected transwells after exposure and stored at −20 °C until analysis. ELISAs for IL-6 and TNF-α (BioTechne) were performed per manufacturer instructions. For each replicate and cytokine, concentrations at baseline (Hour 0) and the later post-exposure time point (Hour 0.25) were paired to compute log2 fold-change (log2FC). A small positive pseudocount equal to half the minimum nonzero concentration across the dataset was added to handle zeros. Log2FC values for each cytokine and condition were tested against 0 (no change) using a one-sample t-test; p-values were BH-adjusted. We report means and 95% CIs of log2FC alongside adjusted p-values; in figures, a broken y-axis is used for clarity when effects are extreme.

### Experimental design and replication

All assays used Caco-2/HT29-MTX-E12 10:1 co-cultures. For resazurin and TEER experiments, conditions included DMEM (control), FMP, PBS/NaCl, diH_2_O, H_2_O, FLEET, SDS, hand soap, and EtOH; replicate numbers per condition were N = 6–9 for TEER and N = 6 for resazurin unless otherwise indicated. For cytokines (IL-6, TNF-α), N = 3 per condition. Where applicable, we performed a preliminary comparison of 10% vs 25% HT29 proportions and observed no significant difference in 48-h viability, so subsequent assays used 10% HT29.

### Statistical analysis

All analyses were performed in R. Data wrangling and summaries used *tidyverse*; specific analyses used *rstatix, emmeans, lme4*, and *splines*. For TEER, we analyzed replicate-level fold-changes relative to baseline, summarized means ± SE over time, and assessed per-time contrasts to DMEM via EMMeans from linear models. For resazurin, we modeled transformed Ab605 (2^Ab605) using linear mixed-effects models with condition-by-time interactions via natural splines and random intercepts for replicates, and we computed marginal means and BH-adjusted pairwise contrasts. For cytokines, we computed per-replicate log2 fold-changes with a strictly positive pseudocount (half the minimum non-zero concentration) and tested mean log2FC against 0 using one-sample t-tests with BH adjustment. Unless noted, two-sided tests were used and BH adjustment controlled the false discovery rate across the family of inferences within each analysis panel. Significance annotations in figures reflect BH-adjusted p-values.

### Data visualization and reporting

Plots were generated with *ggplot2* using *ggbreak* for broken axes and *viridis* vision friendly color palettes. TEER line plots show mean ± SE; heatmaps display mean responses with significance asterisks from BH-adjusted contrasts versus DMEM. Resazurin trajectories show individual replicate traces (faint) with overlaid means; heatmaps annotate BH-adjusted per-time contrasts to DMEM. Cytokine bar plots display means with 95% CIs and BH-adjusted p-value labels. Missing means were imputed by linear interpolation for visualization only (Raw figures in supplemental). Summary tables of final-time endpoints (TEER and resazurin) were produced with gt.

## Contributions

Conceptualization and Design: NRK, EG, ECS. Methodology: NRK and ECS. Data Analysis and Visualization: ECS. Writing, Original Draft: ECS. Writing, Editing: NRK, EG, ECS, CMB, GL. Research Supervision: NRK.

## Research Funding

None.

